## Supplementary figures and images for "A STIM dependent dopamine-insulin axis maintains the larval drive to feed and grow in Drosophila"

### Supplemental Figure 1

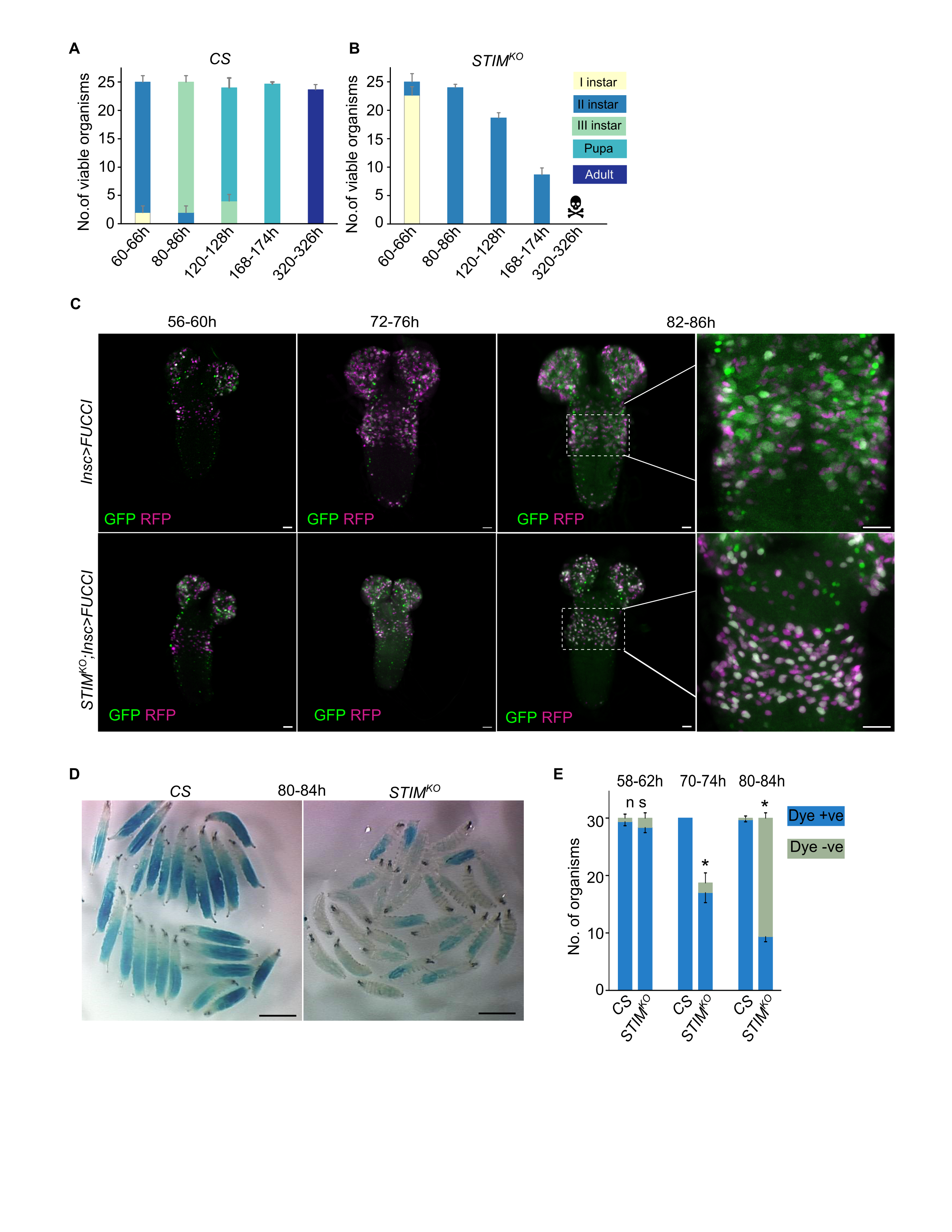

### Supplemental Figure 2

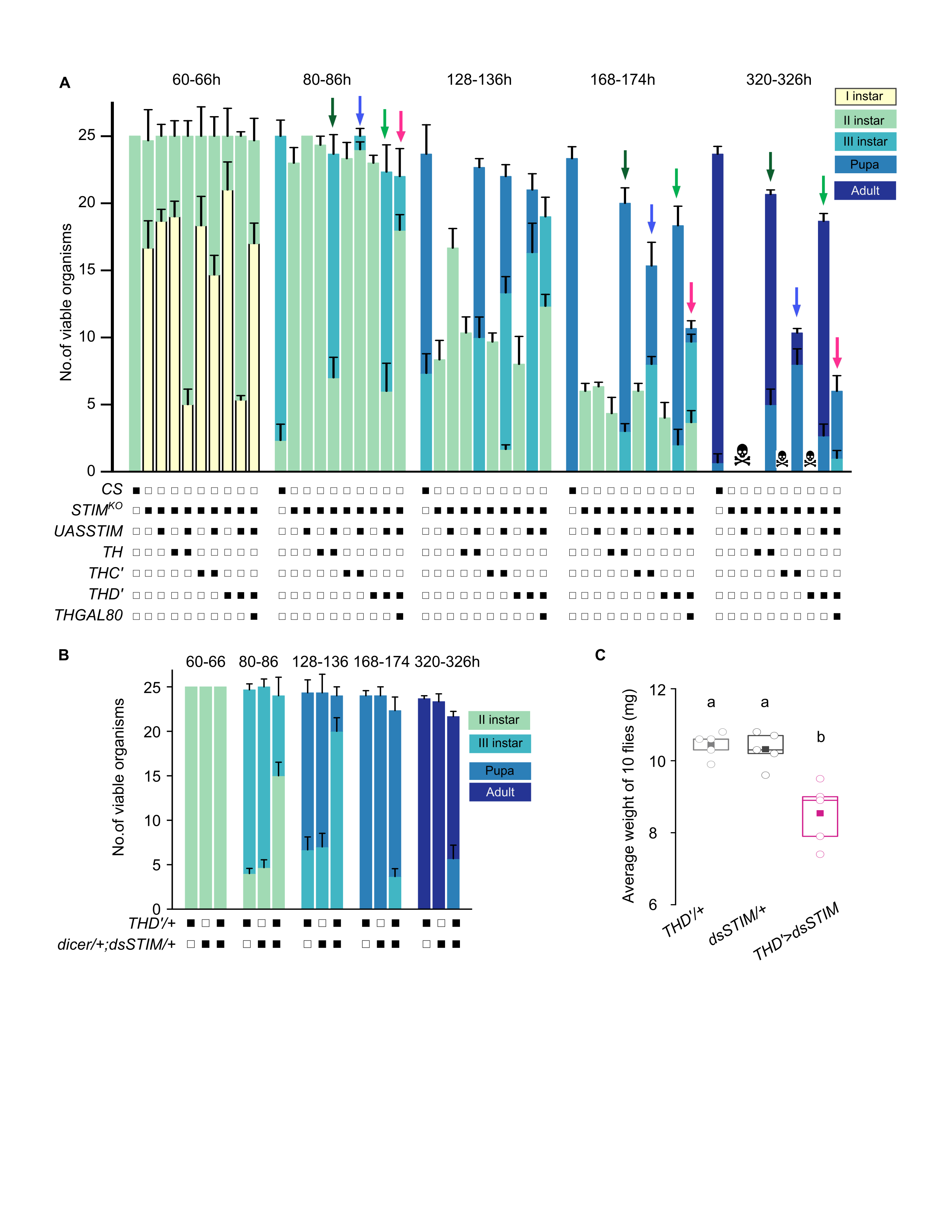

### Supplemental Figure 3

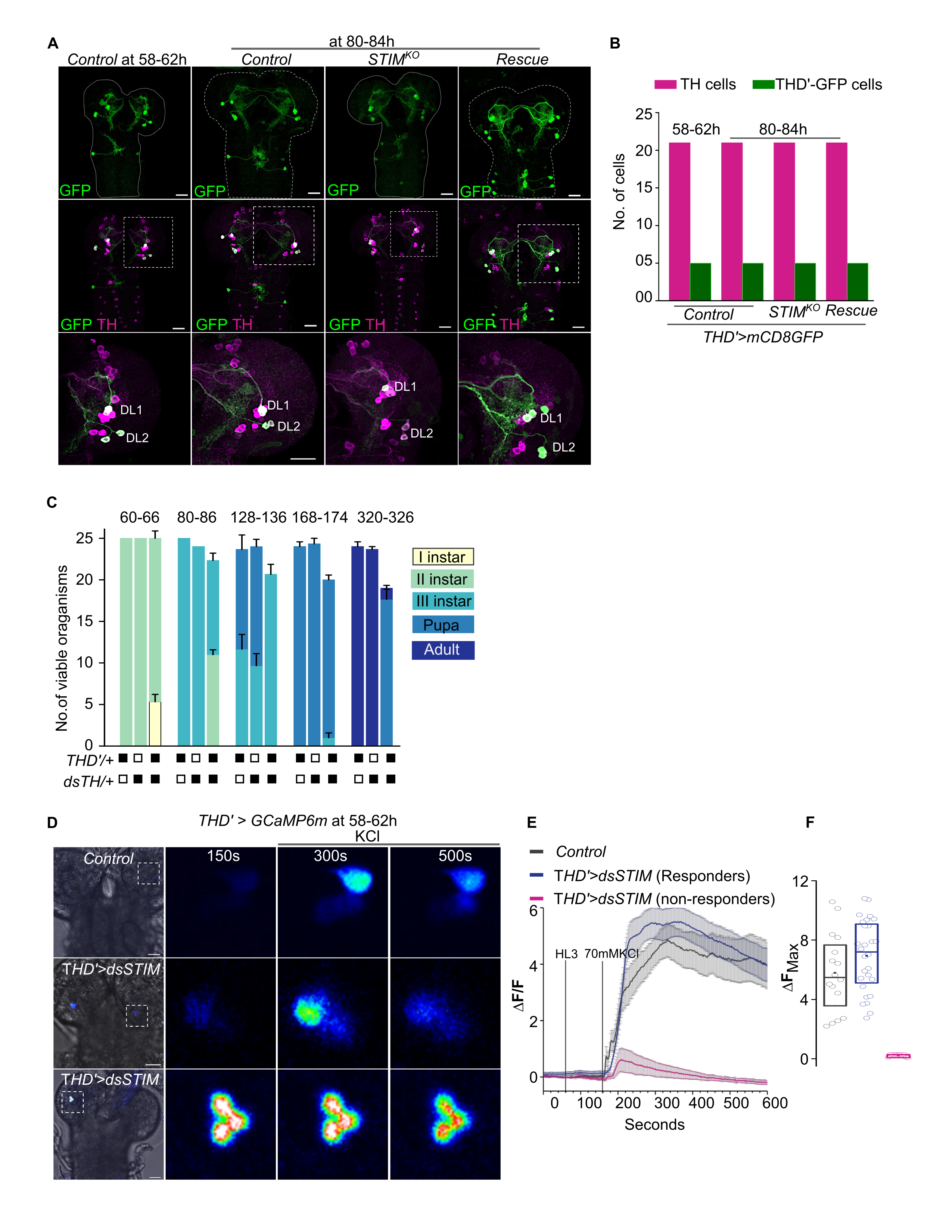

### Supplemental Figure 4

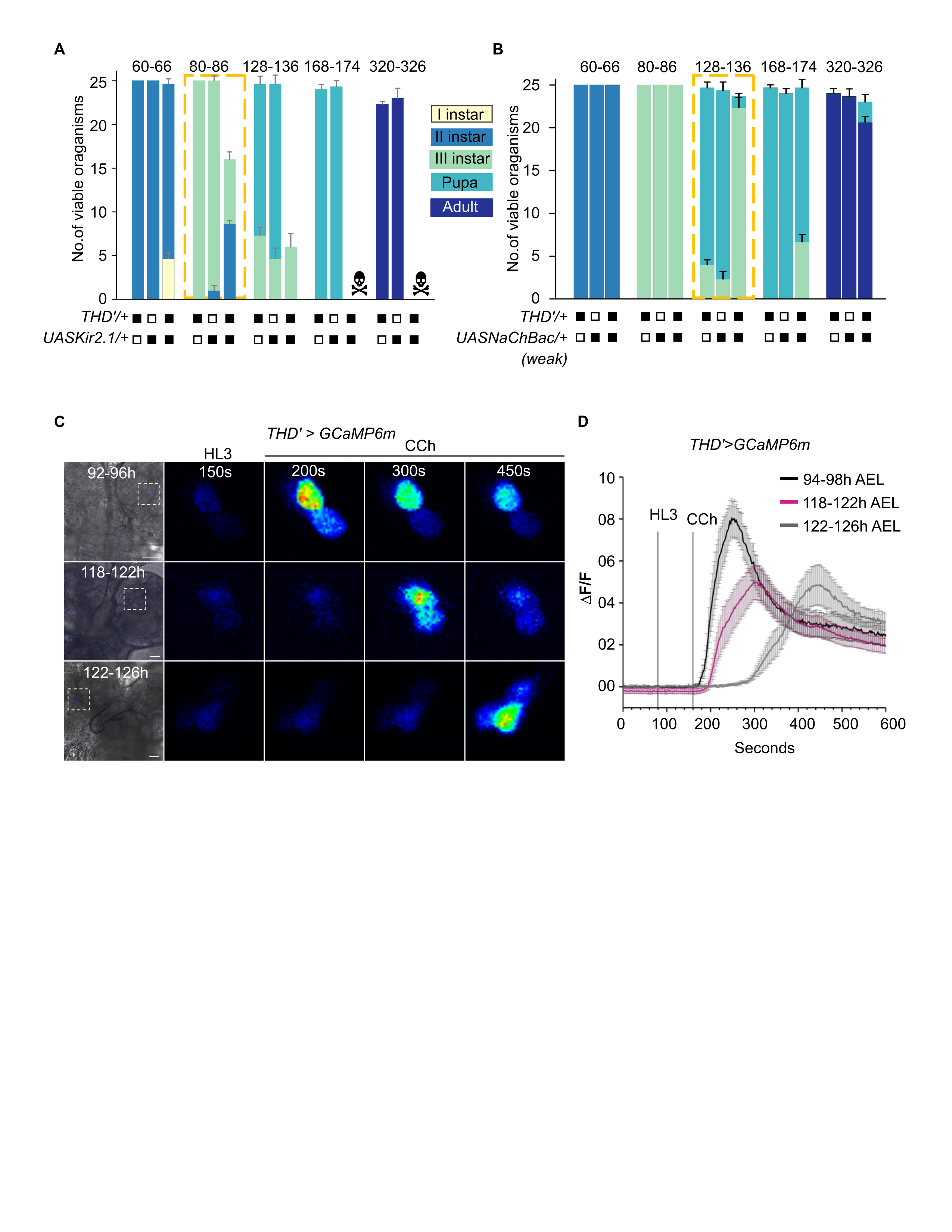

### Supplemental Figure 5

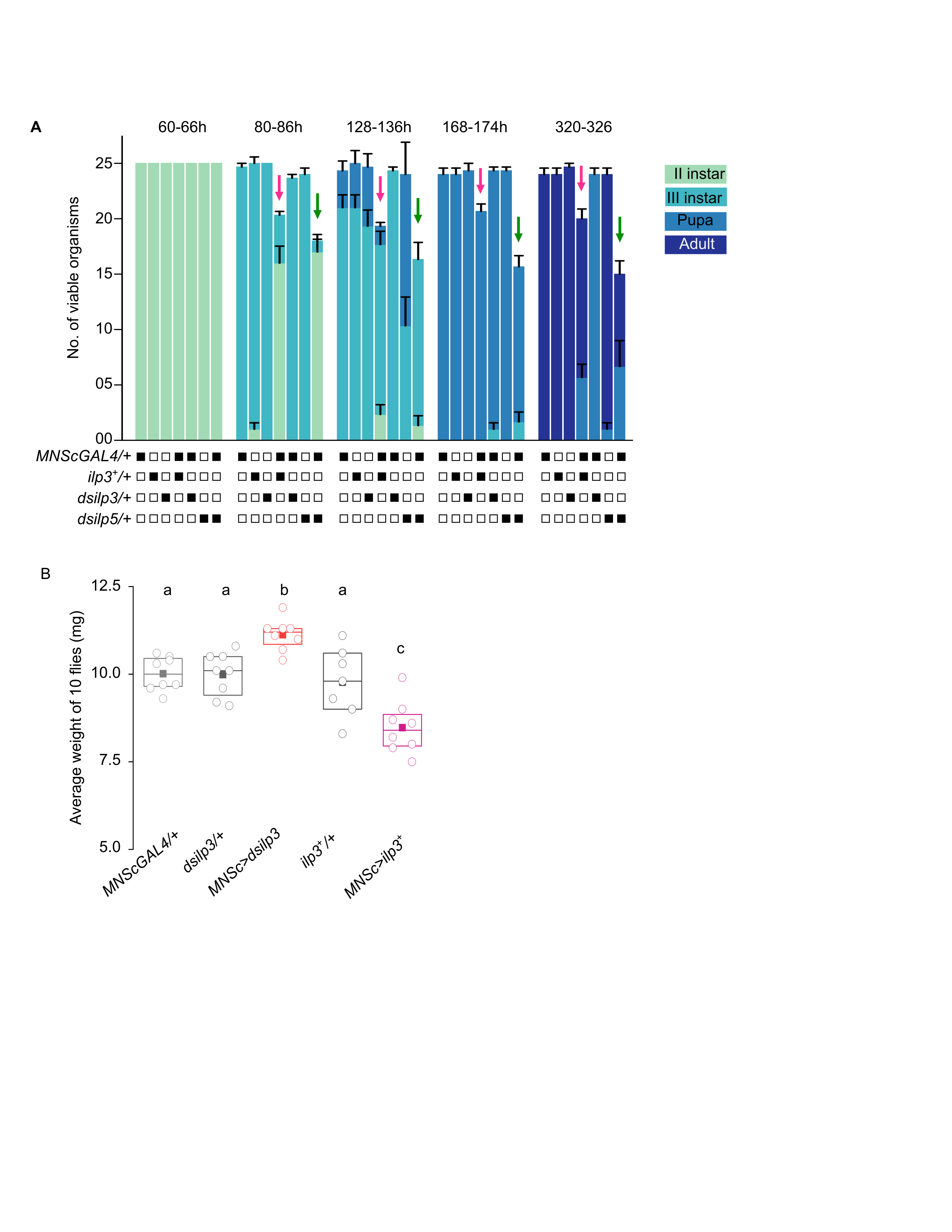
