## Supplemental Table 1 for "A STIM dependent dopamine-insulin axis maintains the larval drive to feed and grow in Drosophila"

| **Gene name** | ***CS*** | ***STIM^KO^*** | **Log2fold** | **p-value** |
| --- | --- | --- | --- | --- |
| *Ilp2* | 278.6493 | 60.52423 | -2.207 | 0.0598 |
| *Ilp3* | 2.516758 | 30.66396 | +3.418 | 0.0032 |
| *Ilp5* | 130.6938 | 7.366336 | -4.166 | 0.0024 |
| *InR* | 9.445050518 | 38.05799706 | +1.951 | 0.0105 |
| *Ilp6* | 28.88770 | 17.25519 | -0.755 | 0.4784 |
| *Ilp7* | 75.89015 | 54.04743 | -0.481 | 0.6072 |
| *CCHa1* | 52.93767 | 173.7131 | +1.686 | 0.0446 |
| *LK* | 462.6413 | 1354.287 | +1.546 | 0.0618 |
| *Sifa* | 116.1298 | 44.68951 | -1.377 | 0.0232 |
| *AstA* | 53.96644 | 93.98807 | +0.831 | 0.1853 |
| *PDF* | 98.60125 | 92.96726 | -0.071 | 0.9157 |
| *CCHa2* | 2.113158 | 7.342901 | +1.599 | 0.1952 |
| *Lst* | 7.543376 | 3.688641 | +0.864 | 0.5004 |
| *Akh* | 145.5645 | 333.8186 | +1.190 | 0.1600 |
| *Crz* | 332.7732 | 576.2431 | +0.787 | 0.2208 |
| *sNPF* | 2867.630 | 5488.131 | +0.936 | 0.1941 |
| *DH44* | 301.0334 | 333.9668 | +0.149 | 0.7875 |
| *DSk* | 53.60334 | 36.63676 | +0.547 | 0.6110 |
| *PTTH* | 1.878419 | 1.306375 | -0.733 | 0.6971 |
| *FMRFa* | 206.0697 | 294.7297 | -0.515 | 0.4000 |
| *ITP* | 23.63023 | 32.28403 | +0.458 | 0.5496 |
| *Rya* | 0.846811 | 1.815414 | +2.264 | 0.3660 |
| *GPA2* | 99.54322 | 183.6279 | +0.887 | 0.3712 |

**Supplementary table 1:** Changes in expression of neuropeptide and receptor encoding genes in  *STIM^KO^* central nervous system
