## Supplemental Table 2 for "A STIM dependent dopamine-insulin axis maintains the larval drive to feed and grow in Drosophila"

**Supplementary Table 2:** P values for main and supplementary figures.

| Figure | Stage of analysis | | Comparison | P value |
| --- | --- | --- | --- | --- |
| 1A | 1^st^ instar | 48h | *CS vs STIM^KO^* | 0.0076 |
| 1B | 2^nd^ instar | 60h | *CS vs STIM^KO^* | 0.0010 |
|  |  | 72h | *CS vs STIM^KO^* | 0.1720 |
|  |  | 76h | *CS vs STIM^KO^* | 0.0395 |
| 1E | 36h | | *CS vs STIM^KO^* | 4.63E-06 |
|  | 44h | | *CS vs STIM^KO^* | 9.22E-05 |
|  | 60h | | *CS vs STIM^KO^* | 3.93E-10 |
|  | 72h | | *CS vs STIM^KO^* | 9.49E-14 |
|  | 84h | | *CS vs STIM^KO^* | 9.24E-14 |
| 1H | 70-74 h | | Dividing neuroblast | 0.0159 |
|  |  |  | Non-dividing neuroblast | 0.0838 |
|  | 82-86 h | | Dividing neuroblast | 0.0026 |
|  |  |  | Non-dividing neuroblast | 0.0043 |
| 1J | 40-44h | | *CS vs STIM^KO^* | 0.0127 |
|  | 58-62h | | *CS vs STIM^KO^* | 0.0270 |
|  | 80-84h | | *CS vs STIM^KO^* | 0.0060 |
| 1K | 36h | | *CS vs STIM^KO^* | 0.0229 |
|  | 42h | | *CS vs STIM^KO^* | 5.51E-07 |
|  | 54h | | *CS vs STIM^KO^* | 0.0004 |
|  | 60h | | *CS vs STIM^KO^* | 1.07E-14 |
|  | 72h | | *CS vs STIM^KO^* | 1.89E-05 |
|  | 84h | | *CS vs STIM^KO^* | 8.22E-05 |
| S1D | 70-74h | | *CS vs STIM^KO^* (Dye ^+ive^) | 0.0086 |
|  | 80-84h | | *CS vs STIM^KO^* (Dye ^+ive^) | 0.0001 |
| 2A | 3^rd^ instar | 78h | *CS vs STIM^KO^; THD’ > STIM^+^* | 0.0127 |
|  |  | 84h | *CS vs STIM^KO^; THD’> STIM^+^* | 0.0198 |
|  |  | 90h | *CS vs STIM^KO^; THD’> STIM^+^* | 0.1127 |
|  |  | 96h | *CS vs STIM^KO^; THD’> STIM^+^* | 0.2113 |
| 2B | Adults | 320-326h | *CS vs STIM^KO^; THD’>STIM^+^* | 0.0005 |
| 2E | 3^rd^  instar | 78h | *CS vs STIM^KO^*; *THD’>STIM^+^* | 0.0040 |
|  |  |  | *CS vs STIM^KO^; THD’,THGAL80 > STIM^+^* | 0.0005 |
|  |  |  | *STIM^KO^; THD’>STIM^+^ vs STIM^KO^; THD’,THGAL80 >STIM^+^* | 0.0106 |
|  |  | 84h | *CS vs STIM^KO^*; *THD’>STIM^+^* | 0.0370 |
|  |  |  | *CS vs STIM^KO^;THD’,THGAL80 > STIM^+^* | 0.0009 |
|  |  |  | *STIM^KO^; THD’ > STIM^+^ vs STIM^KO^; THD’ ,THGAL80 > STIM^+^* | 0.0017 |
|  |  | 90h | *CS vs STIM^KO^*; *THD’>STIM^+^* | 0.0370 |
|  |  |  | *CS vs STIM^KO^; THD’,THGAL80 > STIM^+^* | 0.0033 |
|  |  |  | *STIM^KO^; THD’>STIM^+^ vs STIM^KO^; THD’,THGAL80 >STIM^+^* | 0.0018 |
|  |  | 96h | *CS vs STIM^KO^*; *THD’ > STIM^+^* | 0.11270 |
|  |  |  | *CS vs STIM^KO^; THD’,THGAL80 > STIM^+^* | 0.0131 |
|  |  |  | *STIM^KO^; THD’>STIM^+^ vs STIM^KO^; THD’,THGAL80 > STIM^+^* | 0.0130 |
| 2F | Adult | 320-326h | *CS vs STIM^KO^*; *THD’ >STIM^+^* | 0.00588 |
|  |  |  | *CS vs STIM^KO^; THD’,THGAL80 > STIM^+^* | 0.00031 |
|  |  |  | *STIM^KO^; THD’>STIM^+^ vs STIM^KO^; THD’,THGAL80 >STIM^+^* | 0.0058 |
| 2H | 40-44h | | *CS vs STIM^KO^* | 0.0129 |
|  |  |  | *CS vs STIM^KO^*; *THD’>STIM^+^* | 0.397 |
|  |  |  | *STIM^KO^ vs STIM^KO^; THD’>STIM+* | 0.0026 |
|  | 58-62h | | *CS vs STIM^KO^* | 0.0012 |
|  |  |  | *CS vs STIM^KO^*; *THD’>STIM^+^* | 0.2845 |
|  |  |  | *STIM^KO^ vs STIM^KO^;THD’>STIM^+^* | 0.0041 |
|  | 80-84h | | *CS vs STIM^KO^* | 0.0039 |
|  |  |  | *CS vs STIM^KO^*; *THD’>STIM^+^* | 0.2159 |
|  |  |  | *STIM^KO^ vs STIM^KO^; THD’>STIM+* | 0.0065 |
| 2I | 1^st^ instar | 40-44h | *CS vs STIM^KO^* | 0.0017 |
|  |  |  | *CS vs STIM^KO^*; *THD’>STIM^+^* | 0.2945 |
|  |  |  | *STIM^KO^ vs STIM^KO^*; *THD’>STIM^+^* | 0.0030 |
|  | 2^nd^ instar | 70-74h | *CS vs STIM^KO^* | 3.3854E-08 |
|  |  |  | *CS vs STIM^KO^*; *THD’>STIM^+^* | 0.4371 |
|  |  |  | *STIM^KO^ vs STIM^KO^*; *THD’>STIM^+^* | 1.2909E-08 |
| 2J | 3^rd^ instar | 78h | *THD’/+ vs dsSTIM/+* | 0.2958 |
|  |  |  | *THD’/+ vs THD’>dsSTIM* | 0.0007 |
|  |  |  | *dsSTIM/+ vs THD’>dsSTIM* | 0.0045 |
|  |  | 84h | *THD’/+ vs dsSTIM/+* | n. s. |
|  |  |  | *THD’/+ vs THD’ >dsSTIM* | 0.0150 |
|  |  |  | *dsSTIM/+ vs THD’ >dsSTIM* | 0.0150 |
| 2L | 2^nd^ instar | 58-62h | *THD’/+ vs dsSTIM/+* | 0.4676 |
|  |  |  | *THD’/+ vs THD’>dsSTIM* | 0.0068 |
|  |  |  | *dsSTIM/+ vs THD’> dsSTIM* | 0.0073 |
|  | 3^rd^ instar | 82-86h | *THD’/+ vs dsSTIM/+* | 0.2395 |
|  |  |  | *THD’/+ vs THD’>dsSTIM* | 0.0051 |
|  |  |  | *dsSTIM/+ vs THD’ >dsSTIM* | 0.0006 |
| S2A | 2^nd^ instar | 80-86h | *CS vs STIM^KO^* | 0.0001 |
|  |  |  | *CS vs STIM^KO^; STIM^+^/+* | 0.0013 |
|  |  |  | *CS vs STIM^KO^; TH/+* | 0.0002 |
|  |  |  | *CS vs STIM^KO^; THC'/+* | 0.0001 |
|  |  |  | *CS vs STIM^KO^; THD'/+* | 0.0002 |
|  |  |  | *CS vs STIM^KO^; TH> STIM^+^* | 0.0371 |
|  |  |  | *CS vs STIM^KO^; THC'> STIM^+^* | 0.0002 |
|  |  |  | *CS vs STIM^KO^; THD'> STIM^+^* | 0.0076 |
|  |  |  | *CS vs STIM^KO^; THGAL80,THD' > STIM^+^* | 0.0003 |
|  |  |  | *STIM^KO^ vs STIM^KO^; STIM^+^/+* | 0.1127 |
|  |  |  | *STIM^KO^ vs STIM^KO^; TH/+* | 0.1955 |
|  |  |  | *STIM^KO^ vs STIM^KO^; THC'/+* | 0.4256 |
|  |  |  | *STIM^KO^ vs STIM^KO^; THD'/+* | 0.5 |
|  |  |  | *STIM^KO^ vs STIM^KO^; TH>STIM^+^* | 0.0005 |
|  |  |  | *STIM^KO^ vs STIM^KO^; THC'>STIM^+^* | 0.2475 |
|  |  |  | *STIM^KO^ vs STIM^KO^; THD'>STIM^+^* | 0.0028 |
|  |  |  | *STIM^KO^ vs STIM^KO^ THGAL80,THD' STIM^+^* | 0.0187 |
|  |  |  | *STIM^KO^; TH> STIM^+^ vs STIM^KO^; THC'>STIM^+^* | 0.0009 |
|  |  |  | *STIM^KO^; TH>STIM^+^ vs STIM^KO^; THD'>STIM^+^* | 0.3591 |
|  |  |  | *STIM^KO^; THC’>STIM^+^ vs STIM^KO^; THD'>STIM^+^* | 0.0070 |
|  |  |  | *STIM^KO^; THD’>STIM^+^ vs STIM^KO^; THGAL80,THD'>STIM^+^* | 0.0075 |
|  | 3^rd^ instar | 80-86h | *CS vs STIM^KO^; TH>STIM^+^* | 0.0167 |
|  |  |  | *CS vs STIM^KO^ THC'>STIM^+^* | 0.0002 |
|  |  |  | *CS vs STIM^KO^ THD'>STIM^+^* | 0.0373 |
|  |  |  | *CS vs STIM^KO^; THGAL80,THD'>STIM^+^* | 0.0022 |
|  |  |  | *STIM^KO^; TH>STIM^+^ vs STIM^KO^ THC'>STIM^+^* | 0.0037 |
|  |  |  | *STIM^KO^; TH>STIM^+^ vs STIM^KO^ THD'>STIM^+^* | 0.4500 |
|  |  |  | *STIM^KO^; THC’>STIM^+^ vs STIM^KO^; THD'>STIM^+^* | 0.0075 |
|  |  |  | *STIM^KO^; THD’>STIM^+^ vs STIM^KO^; THGAL80,THD'>STIM^+^* | 0.0066 |
|  | Pupa | 128-134h | *CS vs STIM^KO^; TH>STIM^+^* | 0.1249 |
|  |  |  | *CS vs STIM^KO^ THC'>STIM^+^* | 0.023 |
|  |  |  | *CS vs STIM^KO^ THD'>STIM^+^* | 0.0092 |
|  |  |  | *CS vs STIM^KO^; THGAL80,THD'>STIM^+^* | 0.0087 |
|  |  |  | *STIM^KO^; TH>STIM^+^ vs STIM^KO^ THC'>STIM^+^* | 0.0111 |
|  |  |  | *STIM^KO^; TH>STIM^+^ vs STIM^KO^ THD'>STIM^+^* | 0.00504 |
|  |  |  | *STIM^KO^; THC’>STIM^+^ vs STIM^KO^; THD'>STIM^+^* | 0.0301 |
|  |  |  | *STIM^KO^; THD’>STIM^+^ vs STIM^KO^; THGAL80,THD'>STIM^+^* | 0.0301 |
|  | Pupa | 168 -174h | *CS vs STIM^KO^; TH>STIM^+^* | 0.0060 |
|  |  |  | *CS vs STIM^KO^ THC'>STIM^+^* | 0.0019 |
|  |  |  | *CS vs STIM^KO^ THD'>STIM^+^* | 0.0129 |
|  |  |  | *CS vs STIM^KO^; THGAL80,THD'>STIM^+^* | 0.0001 |
|  |  |  | *STIM^KO^; TH>STIM^+^ vs STIM^KO^ THC'>STIM^+^* | 0.0097 |
|  |  |  | *STIM^KO^; TH>STIM^+^ vs STIM^KO^ THD'>STIM^+^* | 0.3687 |
|  |  |  | *STIM^KO^; THC’>STIM^+^ vs STIM^KO^; THD'>STIM^+^* | 0.0084 |
|  |  |  | *STIM^KO^; THD’>STIM^+^ vs STIM^KO^; THGAL80,THD'>STIM^+^* | 0.0011 |
|  | Adult | 320-326h | *CS vs STIM^KO^; TH>STIM^+^* | 0.0008 |
|  |  |  | *CS vs STIM^KO^ THC'>STIM^+^* | 3.6875E-05 |
|  |  |  | *CS vs STIM^KO^ THD'>STIM^+^* |  |
|  |  |  | *STIM^KO^; TH>STIM^+^ vs STIM^KO^ THC'>STIM^+^* | 4.6487E-06 |
|  |  |  | *STIM^KO^; TH>STIM^+^ vs STIM^KO^ THD'>STIM^+^* | 0.3257 |
|  |  |  | *STIM^KO^; THC’>STIM^+^ vs STIM^KO^; THD'>STIM^+^* | 0.0001 |
|  |  |  | *STIM^KO^; THD’>STIM^+^ vs STIM^KO^; THGAL80,THD'>STIM^+^* | 0.0006 |
| S2B | 3^rd^ instar | 80-86h | *THD’/+ vs dsSTIM* | 0.2108 |
|  |  |  | *dsSTIM/+ vs THD’>dsSTIM* | 0.0076 |
|  |  |  | *THD’/+ vs THD’>dsSTIM* | 0.0066 |
|  | Pupa | 128-134h | *THD’/+ vs dsSTIM* | 0.0023 |
|  |  |  | *dsSTIM/+ vs THD’>dsSTIM* | 0.0353 |
|  |  |  | *THD’/+ vs THD’>dsSTIM* | 0.0010 |
| S2C | Adult body weight | 6-10h post eclosion | *THD’/+ vs dsSTIM* | 0.33193 |
|  |  |  | *dsSTIM/+ vs THD’>dsSTIM* | 0.00081 |
|  |  |  | *THD’/+ vs THD’>dsSTIM* | 0.0012 |
| 3A | 3^rd^ instar | 82-86h | *THD’/+ vs dsTH/+* | 0.3436 |
|  |  |  | *dsTH /+ vs THD’>dsTH* | 3.2714E-06 |
|  |  |  | *THD’/+ vs THD’>dsTH* | 2.6012E-05 |
| 3C | Larval length | 82-86h | *THD’/+ vs dsTH/+* | 0.138206191 |
|  |  |  | *dsTH /+ vs THD’>dsTH* | 2.90527E-13 |
|  |  |  | *THD’/+ vs THD’>dsTH* | 1.28857E-11 |
| 3D | Adult body weight | 6-10h post eclosion | *THD’/+ vs dsTH/+* | 0.1889 |
|  |  |  | *dsTH /+ vs THD’>dsTH* | 3.9117E-06 |
|  |  |  | *THD’/+ vs THD’>dsTH* | 2.6436E-06 |
| 3H | 3^rd^ instar | 84h | *CS vs STIM^KO^; THD’>NaChBac* | *0.0003* |
|  |  |  | *STIM^KO^; NaChBac/+ vs STIM^KO^; THD’>NaChBac* | *0.0163* |
|  | 3^rd^ instar | 96h | *CS vs STIM^KO^;THD’ > NaChBac* | 0.0002 |
|  |  |  | *STIM^KO^; NaChBac/+ vsSTIM^KO^; THD’>NaChBac* | 0.0030 |
| 3J | Larval length | 84h | *CS vs STIM^KO^* | 1.66E-09 |
|  |  |  | *CS vs STIM^KO^*;*THD’* > *NaChBac* | 0.0033 |
|  |  |  | *STIM^KO^ vs STIM^KO^*; *THD’* >*NaChBaC* | 6.11E-07 |
| 3K | Adult | 320-326h | *CS vs STIM^KO^*;*THD’* >*NaChBac* | 9.99E-08 |
| S3C | 3^rd^ instar | 80-86h | *THD’/+ vs dsTH/+* | 0.0133 |
|  |  |  | *dsTH /+ vs THD’>dsTH* | 0.0020 |
|  |  |  | *THD’/+ vs THD’>dsTH* | 0.0022 |
|  | 3^rd^ instar | 128-134h | *THD’/+ vs dsTH/+* | 0.2154 |
|  |  |  | *dsTH /+ vs THD’>dsTH* | 0.0021 |
|  |  |  | *THD’/+ vs THD’>dsTH* | 0.0067 |
|  | Pupa | 168-174h | *THD’/+ vs dsTH/+* | 0.3661 |
|  |  |  | *dsTH /+ vs THD’>dsTH* | 0.0638 |
|  |  |  | *THD’/+ vs THD’>dsTH* | 0.0357 |
|  | Adults | 320-326h | *THD’/+ vs dsTH* | 0.3257 |
|  |  |  | *dsTH/+ vs THD’>dsTH* | 2.7967E-05 |
|  |  |  | *THD’/+ vs THD’>dsTH* | 5.9374E-07 |
| S4A | 3^rd^ instar | 80-86h | *THD’/+ vs Kir2.1/+* | 0.2948 |
|  |  |  | *Kir2.1/+ vs THD’> Kir2.1/+* | 0.0003 |
|  |  |  | *THD’/+ vs THD’> Kir2.1* | 0.0006 |
| S4B | Pupa | 128-134h | *THD’/+ vs NaChBac/+* | 0.1740 |
|  |  |  | *NaChBac /+ vs THD’> NaChBac /+* | 6.28E-05 |
|  |  |  | *THD’/+ vs THD’> NaChBac* | 0.001296 |
|  | Adults | at 320-326h | *THD’/+ vs NaChBac/+* | 0.38627 |
|  |  |  | *NaChBac /+ vs THD’> NaChBac /+* | 0.0001 |
|  |  |  | *THD’/+ vs THD’> NaChBac* | 3.43E-05 |
| 6B  6B | qPCR  qPCR | Ilp2 | *CS vs STIM^KO^* | 0.0074 |
|  |  |  | *CS vs STIM^KO^*; *THD’>STIM^+^* | 0.0074 |
|  |  |  | *STIM^KO^ vs STIM^KO^; THD’>STIM^+^* | 0.1409 |
|  |  | Ilp3 | *CS vs STIM^KO^* | 0.0042 |
|  |  |  | *CS vs STIM^KO^*; *THD’>STIM^+^* | 0.1587 |
|  |  |  | *STIM^KO^ vs STIM^KO^; THD’>STIM^+^* | 0.0081 |
|  |  | Ilp5 | *CS vs STIM^KO^* | 0.0002 |
|  |  |  | *CS vs STIM^KO^*; *THD’>STIM^+^* | 0.1688 |
|  |  |  | *STIM^KO^ vs STIM^KO^; THD’>STIM^+^* | 0.0616 |
| 6D | 2^nd^ instar | 80-86h | *CS* vs *STIM^KO^; MNSc/+* | 4.1423E-05 |
|  |  |  | *CS* vs *STIM^KO^; dsilp3/+* | 4.4218E-05 |
|  |  |  | *CS* vs *STIM^KO^; MNSc>dsilp3* | 0.0015 |
|  |  |  | *STIM^KO^; MNSc/+ vs STIM^KO^; dsilp3/+* | 0.4012 |
|  |  |  | *STIM^KO^; MNSc/+ vs STIM^KO^; MNSc>dsilp3* | 0.0455 |
|  |  |  | *STIM^KO^; dsilp3/+ vs STIM^KO^; MNSc>dsilp3* | 0.0455 |
|  | Adults | 320-326h | *CS* vs *STIM^KO^; MNSc>dsilp3* | 0.0003 |
| 6F | Larval length | 118-122h | *CS vs STIM^KO^; MNSc > dsilp3* | 0.0170 |
| 6H | 3^rd^ instar | 132h | *MNSc /+ vs dsilp3/+* | 0.2113 |
|  |  |  | *dsilp3/+ vs MNSc>dsilp3* | 0.0055 |
|  |  |  | *MNSc /+ vs MNSc>dsilp3* | 0.0017 |
|  |  | 144h | *MNSc /+ vs dsilp3/+* | n.s. |
|  |  |  | *dsilp3/+ vs MNSc>dsilp3* | 0.0175 |
|  |  |  | *MNSc /+ vs MNSc>dsilp3* | 0.0175 |
| 6J | Pupal volume | 144-156h | *MNSc /+ vs dsilp3/+* | 0.1501 |
|  |  |  | *dsilp3/+ vs MNSc>dsilp3* | 0.0008 |
|  |  |  | *MNSc /+ vs MNSc>dsilp3* | 0.0001 |
| 6K | 3^rd^ instar | 72h | *MNSc /+ vs ilp3^+^/+* | 0.1039 |
|  |  |  | *ilp3+/+ vs MNSc>ilp3^+^* | 0.0410 |
|  |  |  | *MNSc/+ vs MNSc>ilp3^+^* | 0.0364 |
|  |  | 84h | *MNSc /+ vs ilp3^+^/+* | n.s. |
|  |  |  | *ilp3+/+ vs MNSc>ilp3^+^* | 0.0025 |
|  |  |  | *MNSc/+ vs MNSc>ilp3^+^* | 0.0025 |
|  |  | 96h | *MNSc /+ vs ilp3^+^/+* | n.s. |
|  |  |  | *ilp3+/+ vs MNSc>ilp3^+^* | 0.0040 |
|  |  |  | *MNSc/+ vs MNSc>ilp3^+^* | 0.0040 |
| 6M | Larval length | 94-98h | *MNSc /+ vs ilp3^+^/+* | 0.4238 |
|  |  |  | *ilp3+/+ vs MNSc>ilp3^+^* | 3.8599E-10 |
|  |  |  | *MNSc/+ vs MNSc>ilp3^+^* | 4.1019E-11 |
| 6N | Mouth hook contractions | 82-86h | *MNSc /+ vs ilp3^+^/+* | 0.3753 |
|  |  |  | *ilp3+/+ vs MNSc>ilp3^+^* | 0.2221 |
|  |  |  | *MNSc/+ vs MNSc>ilp3^+^* | 0.3305 |
| S5A  S5A | 3^rd^ instar | 80-86h | *MNSc /+ vs ilp3^+^/+* | 0.1955 |
|  |  |  | *ilp3+/+ vs MNSc>ilp3^+^* | 4.2774E-05 |
|  |  |  | *MNSc/+ vs MNSc>ilp3^+^* | 0.000130 |
|  |  |  | *MNSc /+ vs dsilp5/+* | 0.1955 |
|  |  |  | *dsilp5/+ vs MNSc > dsilp5* | 2.4576E-05 |
|  |  |  | *MNSc/+ vs MNSc > dsilp5* | 4.7248E-06 |
|  | Adults  Adults | 320-326h  320-326h | *MNSc /+ vs ilp3^+^/+* | 2.13184 |
|  |  |  | *ilp3+/+ vs MNSc>ilp3^+^* | 0.0013 |
|  |  |  | *MNSc/+ vs MNSc>ilp3^+^* | 0.00137 |
|  |  |  | *MNSc /+ vs dsilp5/+* | 0.1439 |
|  |  |  | *dsilp5/+ vs MNSc > dsilp5* | 0.0006 |
|  |  |  | *MNSc/+ vs MNSc > dsilp5* | 0.0008 |
|  |  |  | *MNSc /+ vs dsilp3/+* | 0.19550 |
|  |  |  | *dsilp3/+ vs MNSc>dsilp3* | 0.0002 |
|  |  |  | *MNSc /+ vs MNSc>dsilp3* | 9.1283E-05 |
| S5B | Adult weight | 6-10h post eclosion | *MNSc /+ vs ilp3^+^/+* | 0.2844 |
|  |  |  | *ilp3+/+ vs MNSc>ilp3^+^* | 0.0002 |
|  |  |  | *MNSc/+ vs MNSc>ilp3^+^* | 0.00787 |
|  |  |  | *MNSc /+ vs dsilp3/+* | 0.4653 |
|  |  |  | *dsilp3/+ vs MNSc>dsilp3* | 0.00236 |
|  |  |  | *MNSc /+ vs MNSc>dsilp3* | 0.00232 |
| 7A | 2^nd^ instar | 80-86h | *MNSc /+ vs dsDop1R1/+* | 0.1499 |
|  |  |  | *dsDop1R1/+ vs MNSc>dsDop1R1* | 0.0002 |
|  |  |  | *MNSc/+ vs MNSc>dsDop1R1* | 0.0002 |
|  |  |  | *MNSc /+ vs dsDopEcR /+* | 0.0352 |
|  |  |  | *dsDopEcR /+ vs MNSc>dsDopEcR* | 0.0076 |
|  |  |  | *MNSc /+ vs MNSc > dsDopEcR* | 0.0197 |
|  |  |  | *MNSc /+ vs dsDop2R2/+* | 0.3862 |
|  |  |  | *dsDop2R2/+ vs MNS >dsDop2R2* | 0.5 |
|  |  |  | *MNSc/+ vs MNSc > dsDop2R2* | 0.3862 |
|  | 3^rd^ instar | 80-86h | *MNSc /+ vs dsDop1R1/+* | 0.198489 |
|  |  |  | *dsDop1R1/+ vs MNSc>dsDop1R1* | 9.92E-05 |
|  |  |  | *MNSc/+ vs MNSc>dsDop1R1* | 0.001442 |
|  |  |  | *MNSc /+ vs dsDopEcR /+* | 0.0438 |
|  |  |  | *dsDopEcR /+ vs MNSc>dsDopEcR* | 0.0278 |
|  |  |  | *MNSc /+ vs MNSc > dsDopEcR* | 0.04356 |
|  |  |  | *MNSc /+ vs dsDop2R2/+* | 0.5 |
|  |  |  | *dsDop2R2/+ vs MNS >dsDop2R2* | 0.3257 |
|  |  |  | *MNSc/+ vs MNSc > dsDop2R2* | 0.386277 |
|  | Adults | 320-326h | *MNSc /+ vs dsDop1R1/+* | 0.2129 |
|  |  |  | *dsDop1R1/+ vs MNSc>dsDop1R1* | 0.0007 |
|  |  |  | *MNSc/+ vs MNSc>dsDop1R1* | 0.0007 |
|  |  |  | *MNSc /+ vs dsDopEcR /+* | 0.0861 |
|  |  |  | *dsDopEcR /+ vs MNSc>dsDopEcR* | 0.0007 |
|  |  |  | *MNSc /+ vs MNSc > dsDopEcR* | 0.0007 |
|  |  |  | *MNSc /+ vs dsDop2R2/+* | 0.3890 |
|  |  |  | *dsDop2R2/+ vs MNS >dsDop2R2* | 0.0740 |
|  |  |  | *MNSc/+ vs MNSc > dsDop2R2* | 0.0585 |
| 7C | Adult weight | 6-10h post eclosion | *MNSc /+ vs dsDop1R1/+* | 0.31332 |
|  |  |  | *dsDop1R1/+/+ vs MNSc> dsDop1R1* | 4.3081E-05 |
|  |  |  | *MNSc /+ vs MNSc > dsDop1R1* | 3.371E-05 |
